# Constriction rate modulation can drive cell size control and homeostasis in *C. crescentus*

**DOI:** 10.1101/266817

**Authors:** Ambroise Lambert, Aster Vanhecke, Anna Archetti, Seamus Holden, Felix Schaber, Zachary Pincus, Michael T. Laub, Erin Goley, Suliana Manley

**Affiliations:** Institute of Physics, École Polytechnique Fédérale de Lausanne (EPFL), 1015 Lausanne, Switzerland; Centre for Bacterial Cell Biology, Institute for Cell and Molecular Biosciences, Newcastle University, Newcastle upon Tyne NE2 4AX, UK; Department of Genetics, Washington University in St Louis, St. Louis, MO, USA; Department of Developmental Biology, Washington University in St Louis, St. Louis, MO, USA; Department of Biology, Massachusetts Institute of Technology, Cambridge, MA, USA; Howard Hughes Medical Institute, Massachusetts Institute of Technology, Cambridge, MA, USA; Department of Biological Chemistry, Johns Hopkins University School of Medicine, Baltimore, MD, USA

## Abstract

Rod-shaped bacteria typically grow first via sporadic and dispersed elongation along their lateral walls, then via a combination of zonal elongation and constriction at the division site to form the poles of daughter cells. Although constriction comprises up to half of the cell cycle, its impact on cell size control and homeostasis has rarely been considered. To reveal the roles of cell elongation and constriction in bacterial size regulation during cell division, we captured the shape dynamics of *Caulobacter crescentus* with time-lapse structured illumination microscopy and used molecular markers as cell-cycle landmarks. We perturbed constriction rate using a hyperconstriction mutant or fosfomycin inhibition. We report that constriction rate contributes to both size control and homeostasis, by determining elongation during constriction, and by compensating for variation in pre-constriction elongation on a single-cell basis.

## Introduction

Cell size regulation is observed nearly universally among prokaryotes [1], allowing them to both control their size at birth and to homeostatically maintain it over multiple generations [2]. Cell size control and homeostasis are critical for survival: once too small, cells lack the volume required to host the essential machinery of life [3] or initiate chromosome segregation [4], whereas cells that are too large may suffer limitations in nutrient uptake [5] and distribution [6] because of their reliance on diffusive transport.

Size regulation is the outcome of multiple sequential processes that take place during the cell division cycle. Progression through the cell cycle is marked by several key processes, including chromosome replication, segregation, and division into two daughter cells. These processes occur once per cell cycle in bacteria such as *Caulobacter crescentus* [7], in contrast to rapidly proliferating organisms such as *Escherichia coli* [8] and *Bacillus subtilis* in which cells often have multi-fork replication and which can, following nutrient up-shifts, initiate replication multiple times in a single cell cycle. In *C. crescentus*, differentiation from a swarmer to a stalked cell and the initiation of chromosome replication and segregation mark the transition from cell cycle phase G1 to S. The completion of replication marks the end of S phase. Once DNA segregation is completed, cells finish cytokinesis to form sibling stalked and swarmer cells during G2/M [9].

From the perspective of the cell wall, individual *C. crescentus* cells elongate exponentially throughout the cell cycle, as is typical for rod-shaped bacteria. Their growth is divided into a stage of pure elongation as peptidoglycan is inserted sporadically along the lateral walls, followed by a stage of mixed elongation and constriction in G2/M phase during which peptidoglycan is inserted at mid-cell to build two new poles [10,11]. Each stage governed by largely independent molecular machines. In *B. subtilis*, when cells are on average longer at the onset of constriction, they are also on average longer at division [12,13]. This suggests a simple model for cell size control, by modifying the cell length at which the divisome, the multi-protein complex which guides division, begins to generate constriction. Another possibility is that the rate of constriction is modulated; this was shown to be the case for MatP, which coordinates chromosome segregation and pole construction in *E. coli* [14].

Several checkpoints have been proposed to couple bacterial cell size homeostasis to cell cycle phases. Shortly after entry into S phase, chromosome segregation must initiate before the cytokinetic Z-ring can assemble at mid-cell, coordinated by the gradient-forming FtsZ inhibitor MipZ [15]. In *E. coli*, the initiation of chromosome replication requires a fixed volume per origin of replication, leading to a “sizer” [16]. Chromosome replication in G1/S may take a constant duration, referred to as a “timer” [8]. Putative molecular mechanisms have generally relied on the accumulation of proteins above a threshold, such as an “initiator” triggering replication [17], or excess peptidoglycan (PG) cell wall precursors triggering constriction [18]. A model of the latter case predicts a constant addition of volume per cell cycle, or “adder”. Indeed, an adder has been observed for *C. crescentus* under a wide range of growth conditions [19]. Deviations from a pure adder toward a mixed relative timer and adder have also been reported for stalked cells, observed over many generations and a range of different temperatures [20]. Any model incorporating a sizer or adder will allow smaller cells to increase while larger cells decrease in size over generations until both converge to a size set by the constant of addition [21]. Thus, both provide a clear means for a population to achieve size homeostasis.

Remarkably, although constriction makes up a significant portion of the cell cycle in many bacteria [22], its impact on cell size control and homeostasis has rarely been considered. Intriguingly, budding yeast may use constriction rate to modulate their size in response to changes in growth conditions [23]. However, a single-cell study of the contribution of the constriction stage in bacteria has been challenging, in part due to the diffraction-limited size of the constriction site, and the need for corroboration by divisome markers to unambiguously identify constriction onset. Furthermore, direct measurement of the instantaneous constriction rate has not been possible.

Here, we investigated whether, and how cells adjust their constriction rate to achieve cell size control and homeostasis. We used structured illumination microscopy (SIM) [24] to resolve the constriction site diameter and measure the size of synchronized *C. crescentus* cells as they progressed through their cell cycle. We show that perturbing the constriction rate changes cell size, independent of elongation rate. Furthermore, we found that within a population the onset of constriction and its rate are coordinated: cells that elongate more than average before constriction undergo a more rapid constriction, leading to less elongation during constriction, and vice-versa. This compensation leads to a higher fidelity adder than permitted by onset control alone, allowing *C. crescentus* to better maintain its size in the face of biological noise.

## Results

### Perturbing constriction rate changes cell length

To test the role of constriction, we perturbed its rate pharmacologically and genetically. Fosfomycin inhibits the peptidoglycan (PG) synthesis enzyme MurA [25], which slows PG synthesis and therefore the constriction rate. Additionally, the divisome includes cell wall remodeling enzymes including the late-arriving FtsW and FtsI. Several point mutants of the trans-glycosylase FtsW [26] and its cognate transpeptidase FtsI [27], referred to as FtsW**I*, resulted in a gain-of-function phenotype in *C. crescentus* [28]. It was hypothesized that these mutations maintain the enzymes in their active state, and thereby would increase the constriction rate [28].

We resolved cell shape dynamics during the cell cycle by performing dual-color imaging of the inner membrane and divisome proteins (FtsZ-GFP, FtsW-GFP) with time-lapse SIM (Figure 1A, Movies S1-S3) on a synchronized population of cells. We used automated image analysis to quantify cell shape parameters during the cell cycle (Figure 1B, Figure S1, Methods). The overall cell length relative to the wild-type strain was shorter for FtsW**I* and longer for fosfomycin-treated cells, consistent with previous studies [18,28] (Figures 2A, S2A, B).

**Figure 1:**
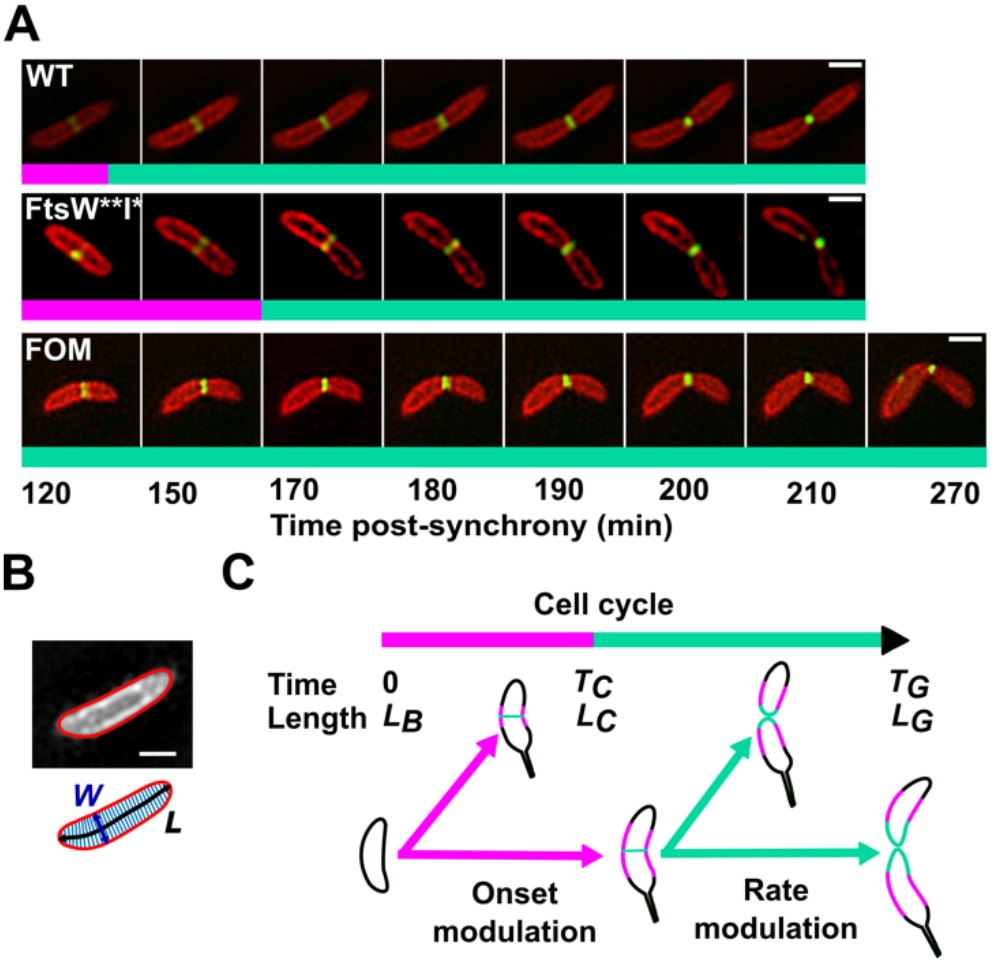
Experimental strategy and constriction-related models for modulation of cell size. (A) Time-lapse SIM images: inner membrane (mCherry-MTS2, red), FtsZ (FtsZ-GFP, green). Shown are example wild-type (WT), FtsW**I* mutant, and fosfomycin-treated cells through constriction, until separation. (B) Analysis of cell shape parameters using sDaDa (see Supplementary methods and Figure S1): the central line (black) is used to measure length (L); the width (W) is extracted from each perpendicular segment; the cell contour defines cell shape (red line). (C) Constriction rate or onset control mechanisms for length. Cells are born at time 0 with length at birth L_B_, and elongate exponentially. T_C_ and L_C_ are the time and length at constriction onset. T_G_ and L_G_ are the time and length at the end of the cell cycle. Magenta parts of the cell contour represent lateral elongation, and cyan parts represent septal elongation. Scale bars: 500 nm. Bicolor bars indicate the stage: pre-constriction (magenta) and post-constriction (cyan).

Could elongation before the onset of constriction (Figure 1C, onset modulation) set the differences in final length between FtsW**I* mutant, fosfomycin-treated and WT cells? The appearance of a measurable constriction in SIM data corresponded well with the arrival of FtsW (Figure S1C), and allowed us to separate elongation before and after constriction onset. Differences in elongation before constriction for all conditions (Figure 2B) were insufficient to account for the observed differences in final length (Figure 2A). Thus, we examined shape changes during constriction (Figure 1C). Individual cells continued to elongate exponentially with the same apparent rate, even as they changed from pure elongation to mid-cell remodeling and constriction (Figures 2 C, E). However, the constriction rate was increased for the FtsW**I* mutant and decreased for fosfomycin-treated cells when compared to WT (Figure 2C), leading to differences in overall cell elongation during constriction (Figure 2D). Thus, we have demonstrated that constriction rate modulation can be a mechanism for cell size control, independent from onset modulation [12,13].

**Figure 2.**
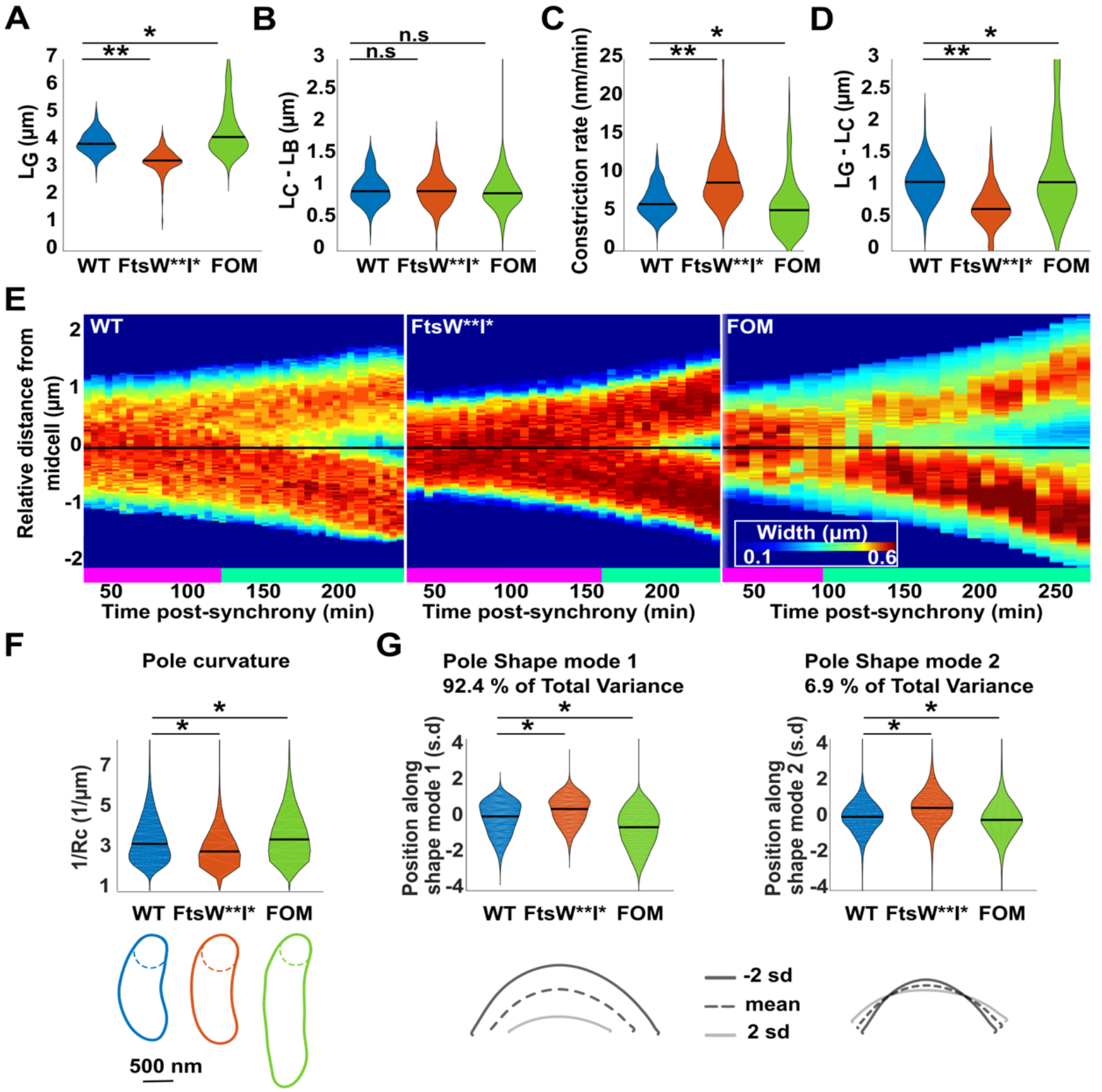
Differences in constriction rate yield different cell sizes and pole shapes. Single-cell distributions of (A) length at division; (B) elongation before constriction; (C) mean constriction rate; (D) elongation during constriction. (A-D), black bars represent the median of the population. Number of cells N: WT: N=208, FtsW**I*: N=212, FOM: N=220. Significance: **: p<0.005, *: p<0.05, n.s.: not significant. (E) Kymographs of representative cells, displaying cell diameter along the cell’s length (vertical axis), versus growth time post-synchrony (horizontal axis), red indicates large diameter, blue small diameter. The middle of the cell is indicated by the black horizontal line. Bicolor bars indicate the stage: pre-constriction (magenta) and post-constriction (cyan). (F) Pole shape analysis. The curvature is the reciprocal of the radius (Rc) of a circle tangent to the curve at a given point, here taken to be the pole. Each cell contour represents a representative single cell from each condition; the distribution of curvatures is plotted above (median value, black bar). (G) The pole region was extracted from each contour (>6000 cells per condition) and analysed using principal component analysis (Celltool, [46]). Shape mode 1 mostly accounts for variation in the length of the pole; shape mode 2 mostly accounts for variation in the bluntness of the pole independent of length. The distributions of each shape mode is plotted, with examples of corresponding shapes, (see also Figure S2)

We found that individual cells continued to elongate at the same rate during constriction, although different perturbations modulated their constriction rate. Thus, faster constriction as in the case of FtsW**I* implies cells should have shorter, blunter poles, while slower constriction as in the case of fosfomycin treatment implies they should have longer, sharper poles. Indeed, kymographs show a more extended gradient in cell width at the poles of fosfomycin-treated cells (Figure 2E). In contrast, FtsW**I* cells show a steeper gradient at the poles. This was confirmed quantitatively by measuring the radius of curvature at the poles (Figure 2F). Furthermore, a population-wide analysis of pole shape demonstrated that over 95% of the total shape variance is accounted for with two principle shape modes, which primarily capture variation in the length and bluntness of the poles (Figure 2G). FtsW**I*, fosfomycin-treated, and WT cells were all distinct along each of these shape axes. We also observed differences in the width of the Z-ring, which appears laterally extended in the fosfomycin case (Figure S2G). This may result from changes in length at constriction onset, since the region of lowest MipZ concentration will be more extended in longer cells [15].

### Elongation before and during constriction compensate one another

To better decipher the relative role of the constriction rate in cell size regulation we further analyzed its contribution to cell size homeostasis. Our experiments were designed to precisely measure the relative contributions to total elongation, and not to distinguish between different general models of homeostasis, which would require measurement over thousands of generations. We found that cells elongated with a distinct mean value for each condition (Figure 3), and that the more individual cells elongated before, the less they elongated during constriction across all conditions tested (Figure 3, Figure S3). Indeed, the total elongation was independent of the relative time cells spent in elongation and constriction phases, with the exception of fosfomycin-treated cells (Figure 3), generally consistent with an “adder.” Consequently, the variance in the total elongation was lower than the variances in elongation before and during constriction would have independently suggested. This was true for all populations, including under perturbed conditions (Figure S3A). These results demonstrate compensation, or over-compensation in the case of fosfomycin (Figure 3, Figure S3B), between elongation before and during constriction resulting in a higher fidelity homeostasis for total elongation (Figure S3A-C).

**Figure 3.**
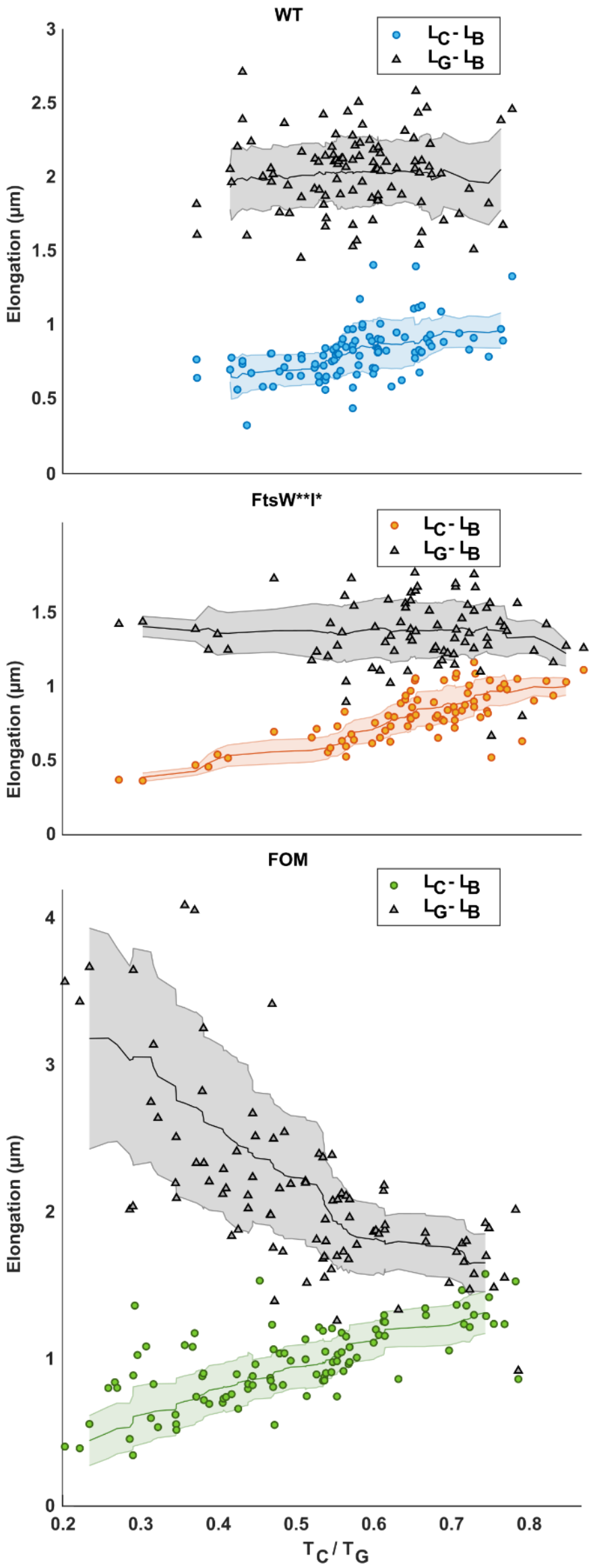
Compensation of elongation before and during constriction contributes to cell size homeostasis. Total elongation (gray) and elongation before constriction (color) for individual Wild Type, FtsW**I* and fosfomycin (FOM)-treated cells, as a function of normalized onset time (T_C_/T_G_). Lines represent the 20 cells moving average; the shaded zones represent the moving standard deviation. Extreme outliers, more than 2 standard deviations from the mean, were omitted for the calculation of the moving average (see also Figure S3).

What could be the mechanism for this compensation? Elongation rate and constriction rate together determine elongation during constriction. Compensation could occur if cells which elongate less before onset subsequently elongate more rapidly or constrict more slowly. However, elongation rate during constriction did not negatively correlate with elongation before constriction (Figure S4A). To better understand constriction dynamics, we examined single cell waist widths as a function of relative duration of constriction (Figure 4A). Cells which elongated more before constriction also spent relatively less time constricting, indicating a higher overall constriction rate. The converse was true for cells which elongated less before constriction, confirming that constriction rate was behind the compensatory effect. Single cells constricted with increasing rate until division; thus, we defined two rates, corresponding to early and late constriction (Figure S4B). Interestingly, early instantaneous constriction rate correlated positively with elongation before constriction, but late rate did not (Figure 4B, C). Hence, early constriction rate changes at the single cell level to adjust elongation during and compensate elongation before constriction.

While molecular mechanisms have been proposed for ensuring homeostasis, the identity of the underlying regulatory factors remains controversial. A previous model estimated PG precursor excess amount as a function of cell cycle [18]. Each cell is assumed to be born with negligible excess, and generates an increasing excess of PG precursors. PG precursors are synthesized in the cell volume, at a volume-dependent rate, while being depleted as they become integrated into the cell wall (Supplementary Note 1). Using this model and experimentally measured cell contours to estimate the changes in surface area (Δ*A*) and volume (Δ*V*), we calculated the excess precursor area (*A*_*excess*_) at the onset of constriction (*T*_*c*_) for individual cells at onset of constriction:

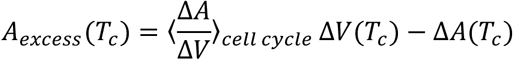

Here, ⟨⟩_*cell cycle*_ refers to the value averaged over the cell cycle. Since it took on average 30 minutes to set up each experiment, we underestimated the volume and area at birth, leading to an offset toward negative estimated precursor excess (Figure 4D). However, we expect the trends to be insensitive to this shift.

**Figure 4.**
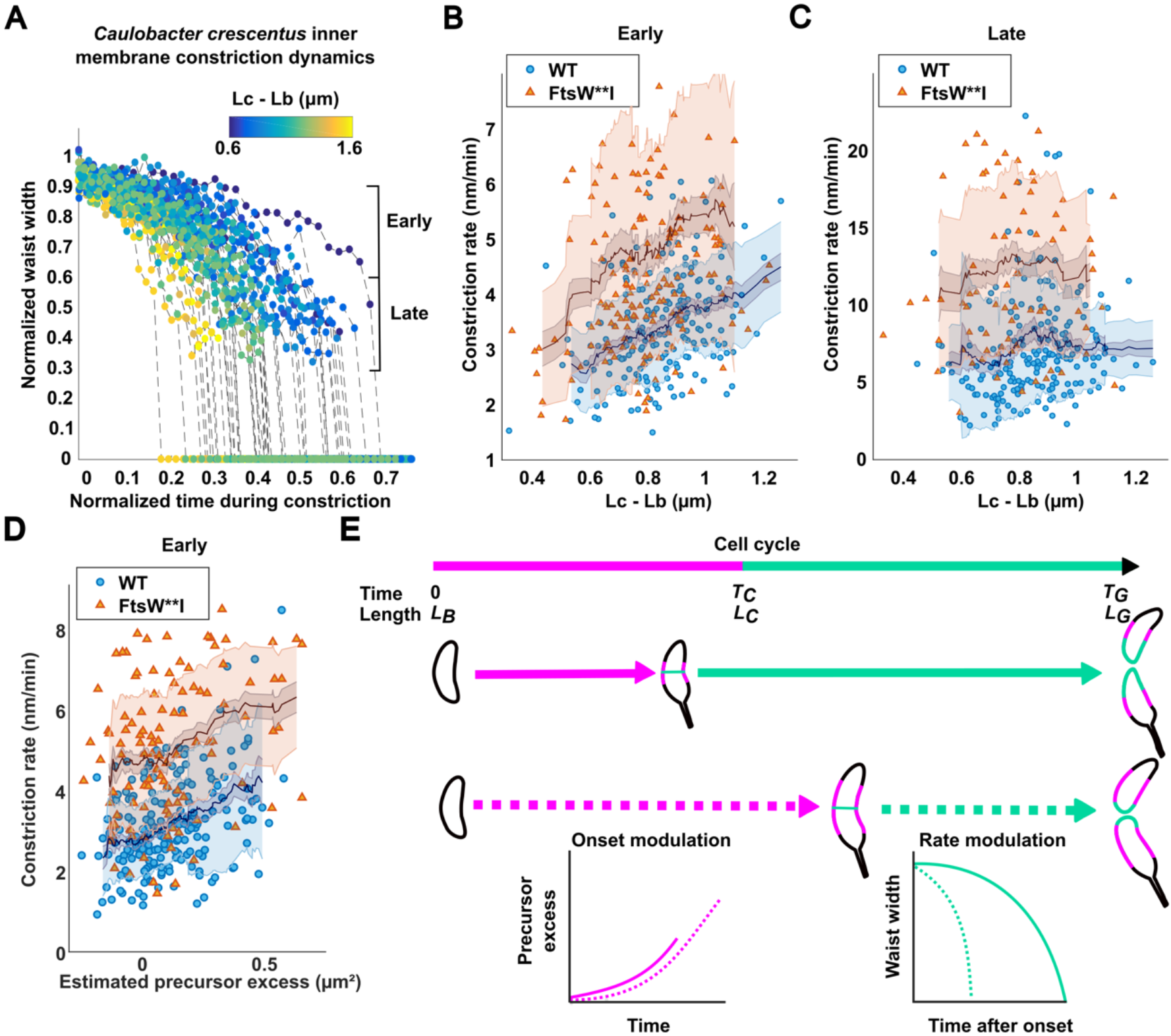
Early constriction rates compensate for elongation before onset. (A) Normalized waist width as a function of normalized time during constriction, colormap represents the elongation before constriction. The measurable constriction was divided into early (0.9-0.6) and late (0.6-0.3) stages. (B) Testing correlation between early constriction rate and elongation before constriction in both WT and mutant strains, WT: r=0.45, p-value<0.01 mutant: r=0.24, p-value<0.01 (Pearson correlation coefficient). (C) Testing correlation between late constriction rate and elongation before constriction, WT: r=0.05, p-value>0.48, mutant: r=0.02, p-value>0.8 (Pearson correlation coefficient). (D) Testing correlation between estimated PG precursor excess and early constriction rate, WT: r=0.33, p-value<0.01, mutant: r=0.29, p-value<0.01 (Pearson correlation coefficient). N>=200 for each strain. (E) Schematic of size regulation in *C. crescentus* with mixed modulation of constriction onset and rate. Magenta parts of the cell contour represent lateral elongation, and cyan parts represent septal elongation. Later onset leads to higher PG precursor excess, which drives more rapid initial constriction (dashed trajectories), and vice-versa (see also Figure S4).

To explain our data, we speculate on a parsimonious model in which PG precursor excess also sets the rate of PG remodeling at the constriction site, and therefore the rate of constriction: the higher the excess the shorter the constriction duration. Indeed, we observeda negative correlation between rate of constriction and estimated excess PG precursor for WT and FtsW**I* cells (Figure 4D, Figure S4). Furthermore, as the new cell poles are built, the excess PG precursor should diminish, leading to a decreased creation of area per time. This is indeed what we observe (supplementary note 2), consistent with models of constriction rate in *E. coli* [14]. Fosfomycin inhibits PG synthesis, so we can no longer use the same mathematical expression to estimate precursor excess, since the activity of fosfomycin would introduce an extra depletion term. Interestingly, within our model this should lead to a slower constriction, consistent with our observations (Figure 2C). Although we have posed the regulatory factor to be PG precursors, this remains controversial because there is only indirect evidence for their role. Any “X-factor” regulatory molecule for constriction rate following the functional relationship described for surface area and volume would fit within the model we suggest.

## Discussion

*Salmonella, E. coli* and *B. subtilis* coordinate their cell size with nutrient availability, perhaps to allow sufficient room for multi-fork replication [4,29,30]. *C. crescentus* shows no such adaptation [19,31], and how its size is modulated has remained a mystery. Our findings show a clear contribution to cell size control from growth during the final constriction stage of cell cycle. Modulation of constriction dynamics changes the overall length of cells, in a manner which has implications for cell shape. In the hypothetical case of extremely rapid constriction, the cell length would be set almost entirely by the growth during the pure elongation stage, leading to short cells with blunt poles. By modulating constriction onset and rate together (Figure 4E), cells may arrive at a variety of pole shapes, an unexpected control mechanism for bacterial cell shape.

Intriguingly, the cell wall itself can have differential properties at the division site. In *B. subtilis*, the division septum contains an enrichment of pentapeptides compared to the rest of the cell envelope [32], perhaps due to a change in the crosslinking or PG composition. In *E. coli*, glycan strands lacking stem peptides are enriched at the septum, allowing proteins containing the PG binding (SPOR) domain to be recruited [33]. In *C. crescentus*, the hydrolase DipM is recruited to the division site by its PG binding LysM domains, suggesting a distinct PG chemistry [32–34]. Consistently, we observed a reduction in staining by wheat germ agglutinin (WGA) at midcell for later stages of the cell (Figure S4I) [37]. Since cells were fixed prior to staining, this implies a difference in the chemistry of the division site, but confounds the interpretation of WGA staining in living cells [20]. We expect that in future studies it will be important to use fluorescent cell-cycle markers in conjunction with fluorescent D-amino acids [11], which together can identify cell-cycle timing and modes of growth. It would be interesting to investigate whether the rate of constriction also affects cell wall chemistry at the division site.

Different factors have been demonstrated to be important for determining constriction dynamics. Before cells can build a septal wall, chromosomes must be partitioned; accordingly, machinery which coordinates the two such as MatP in *E. coli* can also modulate constriction rate [14]. Similarly, dynamically treadmilling FtsZ filaments act as a scaffold to direct cell wall remodelers to the division site, and modulate their rate [38,39]. Nevertheless, the activity of PG remodeling enzymes involved in constriction might depend on PG precursor amounts, and thus may act as a part of a responsive machine. Since, for individual cells, elongation proceeds exponentially with a single rate constant even as PG precursor excess is predicted to increase over cell cycle, the elongation machinery is presumably relatively insensitive to changes in PG precursor amounts. By coupling the elongation machinery to the PG precursor-sensitive constriction machinery, the cell may have arrived at a simple means of compensating for fluctuations in elongation during different phases of the cell cycle. This compensation still has its limitations as we observed in the case of FtsW**I* (Figure S3B); in the case of large elongation before onset, cells must still elongate by a minimum amount during Z-ring maturation (Figure S2H) and constriction. In the future, it will be interesting to identify the molecular partners responsible for constriction rate modulation and PG sensing, and to experimentally investigate the mechanism behind compensation of elongation.

## Acknowledgements

The authors would like to thank Jan Skotheim and Devon Chandler-Brown for critical reading of the manuscript, Manley lab members and Collier lab members for helpful discussions, and the Thanbichler lab for plasmid gifts.

## Author contributions

A.L., A.V., S.M., E.G., M.T.L., S.H. conceived, designed and/or performed experiments, A.L., A.V., A.A., S.M., F. S., S.H., Z.P. created analysis tools and analyzed the data, and all authors contributed to the writing of the manuscript.

## STAR Methods

### CONTACT FOR REAGENT AND RESOURCE SHARING

Further information and requests for resources and reagents should be directed to and will be fulfilled by the Lead Contact, Suliana Manley.

### EXPERIMENTAL MODEL AND SUBJECT DETAILS

#### Bacterial strains and growth conditions

##### Strains and plasmids

The strains, plasmids, oligonucleotides, restriction sites and modes of constructions used for this study are summarized in key resources table. The WT and mutant strains were electroporated with the P_van_ mCherry-MTS2 plasmid to yield the P_van_ mCherry-MTS2 strain. These strains were then electroporated with the P_xyl_ FtsZ-GFP, P_xyl_ FtsW-GFP or P_xyl_ FtsW**-GFP plasmid to yield the respective dual color strains.

##### Growth conditions

Liquid cultures were grown overnight at 28°C with 15 mL of M2G minimal media under mechanical agitation (180 rpm). Each specific inducer for every different condition is described below. Fosfomycin perturbation was achieved with a subminimal inhibitory concentration of 12.5 μg/ml added one hour prior synchronization. To induce the expression of mCherry-MTS2 from the Pvan promoter, 0.5 mM of vanillate was added to the culture before overnight growth. For the expression of FtsW-eGFP, xylose was added to reach a final concentration of 0.3 % (mass per volume) 2 hours before synchrony as optimized previously [40]. P_xyl_ FtsZ-eGFP was induced overnight at 0.003 % xylose in M2G as optimized in a previous study [41].

## METHOD DETAILS

### Sample preparation

Cells were synchronized at 4°C by Percoll density gradient [42] when they reached mid-exponential phase (0D_660_=0.3-0.5). A silicone gasket (Grace Biolabs, 103280) was placed on a rectangular cover slide, and filled with 1% M2G agarose (Ultra PureTM Agarose, Sigma) containing fosfomycin, xylose and vanillate at the appropriate concentrations when needed. Vanillate was present at 0.5 mM in all experiments, xylose was present at 0.003% for induction of FtsZ-eGFP, but absent for the FtsW-eGFP experiments. Fosfomycin was added to a final concentration of 10 μg/ml for drug perturbation experiments. A cover slide was placed on top of the silicone gasket before solidification of the agarose to achieve a flat agarose pad. After 5 min, the top cover slide was removed, and a 1 μL drop of a synchronized cells suspension was placed on the pad. A small piece of agarose (~1 mm) was cut out on two opposing sides to ensure aerobic conditions during imaging. After absorption of the droplet, the pad was sealed with a plasma-cleaned #1.5 round coverslip with a diameter of 25 mm.

### Image acquisition

#### Microscope set up

SIM microscopy was performed on the 3D NSIM Nikon microscope, with a CFI Apochromat TIRF objective (100 x, NA 1.49, Nikon). The microscope was equipped with 400 mW, 561 nm and 480mW, 488 nm lasers (Coherent Sapphire) and a back-illuminated EMCCD camera (iXon 3, Andor Technology) with a 512×512 pixel CCD sensor.

#### Acquisition settings

Dual color imaging of the cells was performed at 28°C using the 488 nm and 561 nm lasers for the divisome protein-eGFP channel and the mCherry-MTS2 channel respectively. The camera was operated with a readout speed of 1 MHz and a dynamic range of 16 bit to have the maximum pixel readout speed at the highest dynamic range. The preamplifier gain and the electron multiplication gain were set to 1 and 200 respectively to maximize the signal to noise at the chosen dynamic range. All raw SIM images were acquired with a camera acquisition time of 200 ms and 100 ms (5 fps and 10 fps) for the 561 and 488 channels. The laser power for both channels was 4 W/cm2. These settings yielded a good balance between image quality and photo-bleaching.

All the raw images were acquired in 3D SIM image mode to ensure the highest signal to noise ratio and lateral resolution. Fifteen images were captured of each 30.7×30.7 μm field of view, five phase-shifted images per angle at each of three interference pattern angles. A full raw dual color image stack was acquired in 17s.

Live-cell fluorescence microscopy over the cell cycle was achieved by performing time-lapse imaging. 3D SIM snapshots were captured at 5 min or 10 min (for fosfomycin-treated cells) time intervals to follow dynamics while minimizing photo-bleaching of the sample during the image acquisition. Multiple fields of view were imaged sequentially at each time point, allowing following up to 200 cells per experiment. Super-resolved SIM images were reconstructed by the Nikon NIS-Elements software.

## QUANTIFICATION AND STATISTICAL ANALYSIS

### Analysis of cell shape dynamics

The super-resolved SIM images were processed via a custom-made software package called sDADA (Shape Dynamics Automated Data Analysis). sDADA generates scatter plots, histograms and violin plots in order to study key parameters controlling the cell size and homeostasis, such as: elongation rate, constriction duration, length at birth, onset, onset time. sDADA extracts these parameters from the analysis of the cell shape dynamics thanks to semi-interactive modules for image segmentation, edge detection, cell filtering, cell tracking and statistical analysis (Supplementary Methods). The MATLAB-based software package is available together with its documentation upon request.

### Parameter definition

We assumed as time zero (T_0_) the time at which the suspension of synchronized bacteria is added to the agarose pad. This occurred approximately 20-40 minutes before starting time-lapse SIM acquisition. T_c_, T_z_, and T_w_ refer to the constriction onset time measured with different approaches. T_z_ is the time of the FtsZ assembly, which we assumed to occur when the fluorescence intensity of FtsZ-eGFP at midcell was three times higher than elsewhere. T_w_ is the FtsW arrival time measured as the moment at which the FtsW signal appeared stable at midcell (Figure S1C).

T_C_ is defined as the time at which the constriction invagination depth is equal to a predetermined normalized waist width threshold. To find the optimal threshold, we tested different thresholds in the reasonable range from 80% up to 99% of the maximum diameter, with step sizes of 1 %. Since FtsW arrival time is an alternative readout of the constriction start the T_C_ values computed from the waist diameter versus time should strongly and robustly correlate with the T_W_ values. For each threshold, we computed the Pearson’s correlation coefficient for the scatter plot of the T_C_ values versus the Tw values of all the cells. We found that a threshold of 92% had the best correlation coefficient and minimal least square error for T_C_ versus T_W_.

Generation time (T_G_) and final length (L_G_) are the time and the length at which the cell divide. Constriction duration τ is the difference between T_G_ and T_C_, or when specified, T_W_. The length at birth, L_B_ and the elongation rate k are extracted from the exponential fitting of the elongation: *L*(*t*) = *L*_B_ · *e*^*kt*^.The length at the constriction onset, L_C_, L_Z_ or L_W_, were measured as the lengths at T_C_, T_Z_ and T_Z_ respectively. The total elongation is the difference between L_G_ and L_B_. Elongation before constriction and during constriction are defined respectively as the difference between L_G_ and L_C_, and between L_C_ and L_B_. When specified, L_w_ can be used instead of L_C_.

### Statistical tests

For each parameter defined in the section above, the statistical significance of observed differences between strains or conditions was tested. We compared the means of the repeats using Mood’s median test (Supplementary Methods). The correlation between variables was analyzed using Pearson’s correlation coefficient in the presence of a linear trend or the Spearman’s correlation coefficient for nonlinear relationship. The experiments were performed in two independent replicates per condition with a minimum of 100 cells analyzed per replicate, and 200 minimum per condition.

## DATA AND SOFTWARE AVAILABILITY

All data and software used to support the results of this manuscript are available from the Lead Contact upon reasonable request. Original data is available on Zenodo, DOI: 10.5281/zenodo.1172042 and 10.5281/zenodo.1172072

## KEY RESOURCES TABLE

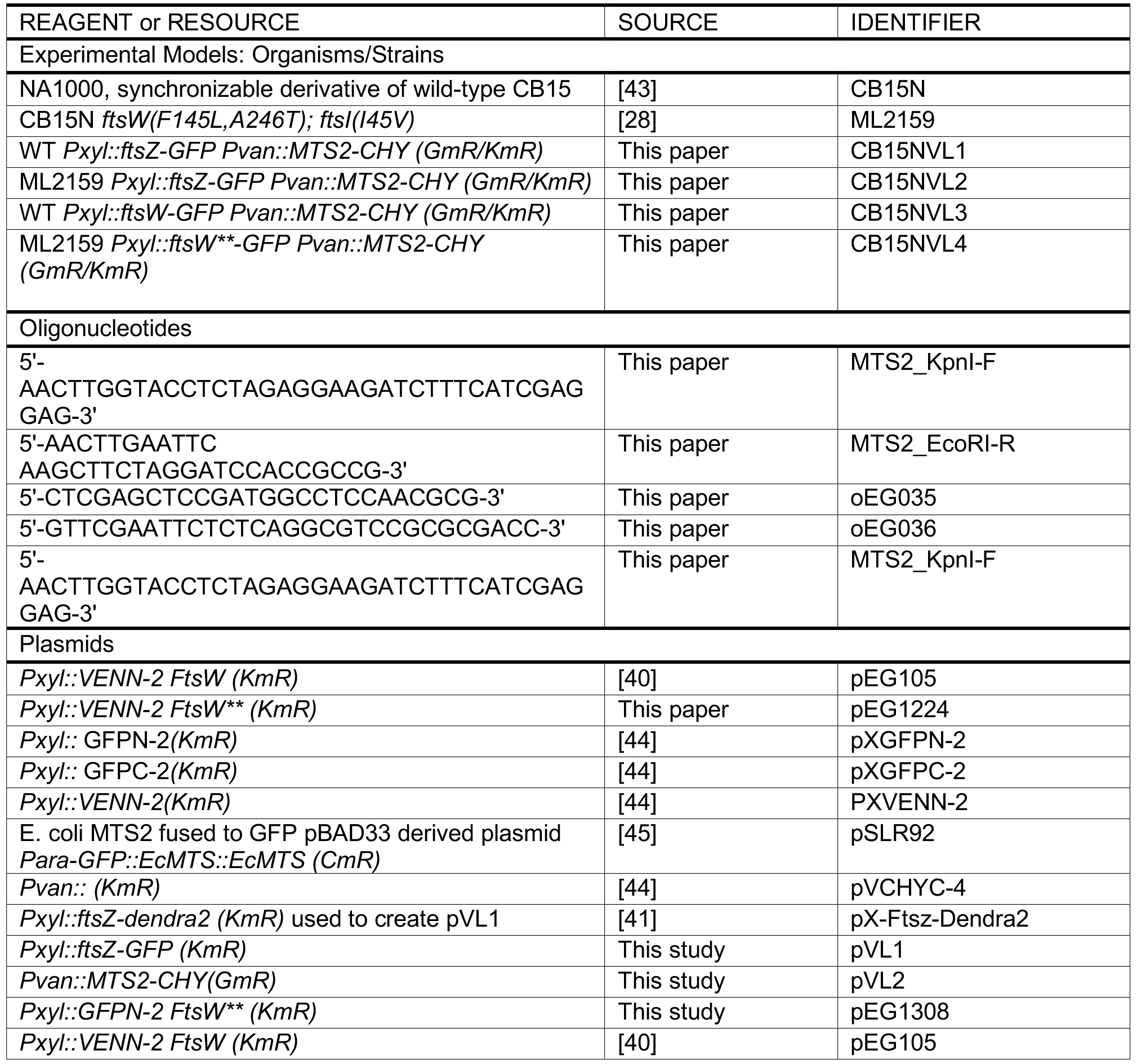

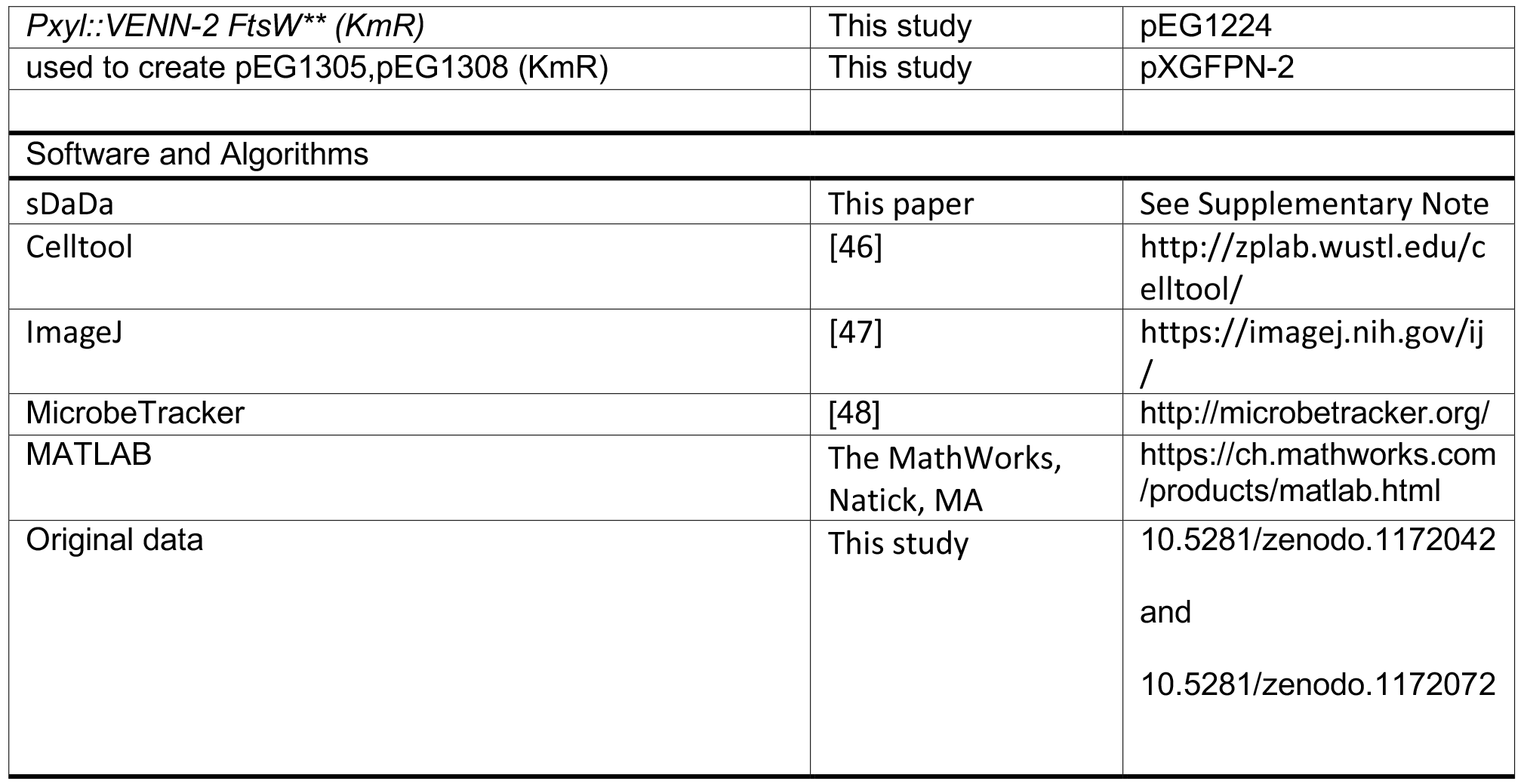

## Supplementary Methods. Related to Figure 1. and methods. Image analysis workflow. Cell shape dynamics and fluorescence measurement

A robust semi-automated pipeline was developed to identify, track and measure shape parameters (length, width, waist width) of hundreds of cells imaged in time lapse movies over the all cell cycle. Also distributions of the variables and correlations between variables were analyzed using this pipeline.

### Drift correction

To simplify the tracking of cells over time, the time-lapse videos were drift-corrected using the Fiji plugin “Descriptor based registration (2d/3d + t)” [1]. The dual-color experiment, registration of both channels was performed based on the drift observed in the red-channel (561nm).

### sDaDa: a software package for supervised segmentation and measurement of bacterial images

The quantitative image analysis of the cell shape dynamics and of their fluorescent signals was performed with our custom-made software package called sDaDa (Shape Dynamics Automated Data Analysis). sDaDa is an open source MATLAB-based program for time-lapse dual color images of bacteria (FtsZ-GFP, FtsW-GFP (green channel) and mCherry-MTS2 [2] (red channel)).

The program takes as input the time-lapse images stack (each superresolved image with a size of 1024 pixel × 1024 pixel; 30 nm /pixel), the camera parameter (i.e. pixel size) and a set of experimental parameters (i.e. starting time, time interval between two consecutive frames, the field of view (FOV)). The program outputs a data structure containing all the measurements of each cell at each frame and a set of figures with the results of the measurements.

The description of the main features of the sDaDa program and of the general steps in a typical pipeline (Figure S1) are presented below.

The main pipeline stages of the software are (1) image segmentation and first edge detection guess, (2) cell tracking, (3) edge detection refinement and shape parameters measurement, (4) divisome ring identification (5) shape parameters correlation analysis and outputs display.

More specifically, the segmentation step (1) is based on two processes: the first is based on the Otsu’s thresholding method [3] to distinguish background pixels from foreground one; the second group together a set of pixels by seeing which pixels are connected to each other. The Otsu’s method sets a proper threshold via maximizing the inter-class variance of the bi-modal histogram of the pixel intensities (foreground pixels and background pixels). Connected pixels that have a signal value higher than the threshold are tagged with the same number and identify as part of a cell only if they form an area bigger than 0.5 μm2. We used two built-in MATLAB functions (graythreshold() and bwlabel()) for Otsu’s image thresholding and to identify each individual cell.

A microbeTracker function named model2mesh.m defines the first cell contour guess starting from the edge detection performed with buit-in MATLAB functions bwperim()and bwtraceboundary()). The model2mesh function retours two semi-contours, corresponding to the ‘left’ and ‘right’ sides of the cell. From the two semi-contours, the bacteria poles (the two farthest apart points on the contour) and the centerline (the average of the two half contour parts) can be easily identified.

Within the segmentation stage, the cell shape search, control and refinement is done thanks to an interactive tool.The user has several tools inspired by the MicrobeTracker [4] approach to manipulate the region, such as removing parts, joining two regions, smoothing, expanding. The user can also choose to delete the current time point of the cell or to mark the cell as divided, after which it will no longer be followed.

In order to track the same bacteria in successive frames (step (2)), the program performs a search based on a spatial analysis. The area and the barycenter position belonging to a cell in one frame is compared with the spatial distribution of pixels belonging to the possible corresponding cell in the following frame. The two regions correspond to the same cell if the difference between these two spatial distributions is lower than a user-defined tolerance parameter. The tracking search stops when the cell divides or when one region does not pass the search criteria. The FtsZ and FtsW divisome assembly time (T_W_ and T_Z_) is determined by monitoring the intensity profile along the centerline length over the cell cycle (step (4)). T_W_ and T_Z_ are the moments at which FtsZ and FtsW fluorescence signals reach their maximum intensity at the midcell.

The segmented regions, containing a well-identified cell, can therefore enter the second stage (3) where the program extracts and accurately measures the shape parameters described below. The diameter is measured by taking perpendicular slices of the bacteria image along its centerline length. Using the intensity profile along each slice, a histogram with two maxima corresponding to the cell edge will define the diameter. The intersection of the histogram with a line parallel to the abscise axis at half maximum high, identify up to four abscises (two for each maximum). The diameter is the difference between the two furthest apart abscises. Repeating this procedure for each slice of the bacteria, the diameter profile as a function of the length could be achieved (Supplementary Fig. 2). The minimum between two maxima of the diameter as a function of the length will then define the measure and the position of the constriction site. As a consequence, the waist width will be easily defined: the ratio of the width of the constriction site and the maximum diameter along the cell. Lastly, the length is measured by calculating the arc length of the centerline.

The program examines the temporal evolution of the length and the waist width for all single cells detected in a time-lapse experiment. From these curves, it extracts the elongation rate, the length at birth (L_B_) and the division time (T_G_). The onset time TC and the duration of the constriction are measured as explained in the Methods “Parameter definition” section.

The volume and the surface area of a cell are estimated based on the measured widths along the length of the bacterium. The width versus the length profile is first smoothed using a spline function, to filter out the noise that would inflate the surface area estimation. The bacterium is assumed symmetric along its central axis. Therefore, assuming cross-sections perpendicular to the axis are circular, the volume and surface area can be computed by treating the measured ‘segments’ of the bacterium as a series of conical frusta.

The last stage (5) provides a set of statistical tools to study the correlations between the parameters extracted and to measure their average and variance. The user can generate a scatter plot for each possible couple of parameters combination (e.g. the scatter plots in Figure 3). Moreover, the user can generate a violin plot for each parameter to inspect its statistical distribution over the entire population (e.g. the violin plots in Figure 2). To conclude the program computes the correlation coefficient of each couple of parameters.

Analysis of cell poles was performed using the Celltool software package ([5]; http://zplab.wustl.edu/celltool/). Image-derived cell shapes were converted into parametric spline curves [5], and centerlines were fit to each cell shape (as described in [6]), again using Celltool. Cell poles were defined as the position on the cell outline closest to the ends of the centerline, and the curvature at that position was calculated from the first and second derivatives of the parametric spline x(t), y(t): curvature = (x′y″ − y′x″)/( x^′2^ + y^′2^)^3/2^, where prime and double-prime represent the first and second derivatives, respectively. The “pole regions” used for PCA shape analysis were defined as all points within distance *d* from each endpoint, where *d* was set to 5% of the total cell perimeter (so 20% of the cell boundary was counted as one pole or the other). Principle modes of pole-shape variation were computed with Celltool, as previously described [5].

## Supplementary Note 1. Related to Equation 1. Estimation of excess peptidoglycan precursor

The assumptions and derivation are based on the work of Harris and Theriot [7]

### Assumption 1

Peptidoglycan precursor (P) production is proportional to the cell volume (V).

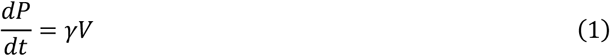

With γ being the rate constant of P production per unit of V, we assume *γ* is constant over the cell cycle (it changes over larger timescales than the generation time).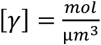

### Assumption 2

PG precursor consumption is proportional to the increase in cell surface area (A):

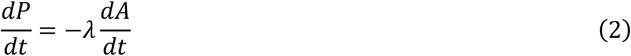

With *λ* being the rate constant of P consumption per unit of A, we assume λ is constant over the cell cycle (it changes over larger timescales than the generation time). 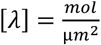

The total rate of change in precursor is then:

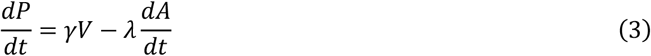

The amount of PG precursors produced between cell birth and an arbitrary time in the cell cycle, *t*_*x*_, is calculated as follows, using exponential Volume growth:

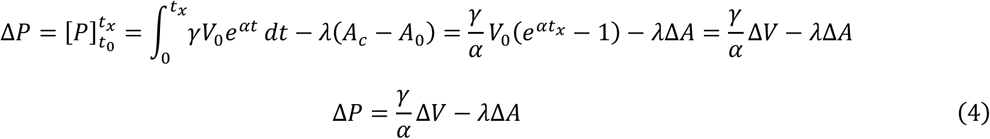

Alternatively:

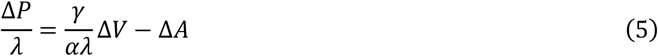

With Δ*P*, Δ*V* and Δ*A* being the increase in P, V and A respectively. 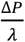 is the excess precursor expressed as the surface area that could be built with it. 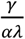 Expresses how volume growth results in production capacity of surface area. To find its value, we can use a **third assumption:** over a cell cycle, between birth and division of a cell, the amount of precursor that is produced equals the amount that is used, or the net production of precursor is zero:

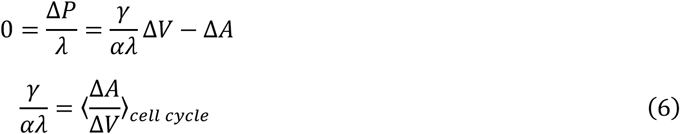

If we apply (6) to (5), we get:

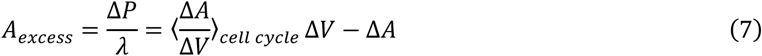

At the onset of constriction, when *t* = *T*_*c*_, this can be written as:

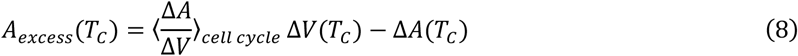

Note that 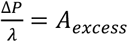 describes the excess precursor as the amount of surface area that could be built with it.

## Supplementary Note 2. Related to Figure S4. Empirical Constriction model

To access we used an empirical model for constriction rate [8]:

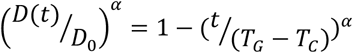

Where D(t) is the diameter of the constriction site in function of time, t, while D_0_ is the diameter at constriction onset, *D*(*t*)/*D*_0_ is the normalized waist width. T_c_ is the time at constriction onset, while T_G_ is the time when constriction and the cell cycle finishes. T_G_-T_C_ is the duration of constriction. α is a variable reflecting the change in constriction rate. For constant constriction rate α equals 1, for a constant buildup of the area of hemispherical poles, α equals 2. Average values for α were 1.4 for the WT and 1.5 FtsW**|, suggesting cell wall remodeling rate slows down.

## Supplementary Movies

**Movie S1. Representative time lapse SIM of *C. crescentus* wild-type.**

Inner membrane (mCherry-MTS2, red), FtsZ (FtsZ-GFP, green), replayed at 10 frames per sec, one frame was recorded every 5 min.

**Movie S2. Representative time lapse SIM of *C. crescentus* FtsW**I*.**

Inner membrane (mCherry-MTS2, red), FtsZ (FtsZ-GFP, green), replayed at 10 frames per sec, one frame was recorded every 5 min.

**Movie S3. Representative time lapse SIM of fosfomycin-treated *C. crescentus*.**

Inner membrane (mCherry-MTS2, red), FtsZ (FtsZ-GFP, green), replayed at 5 frames per sec, one frame was recorded every 10 min.

**Figure S1.**
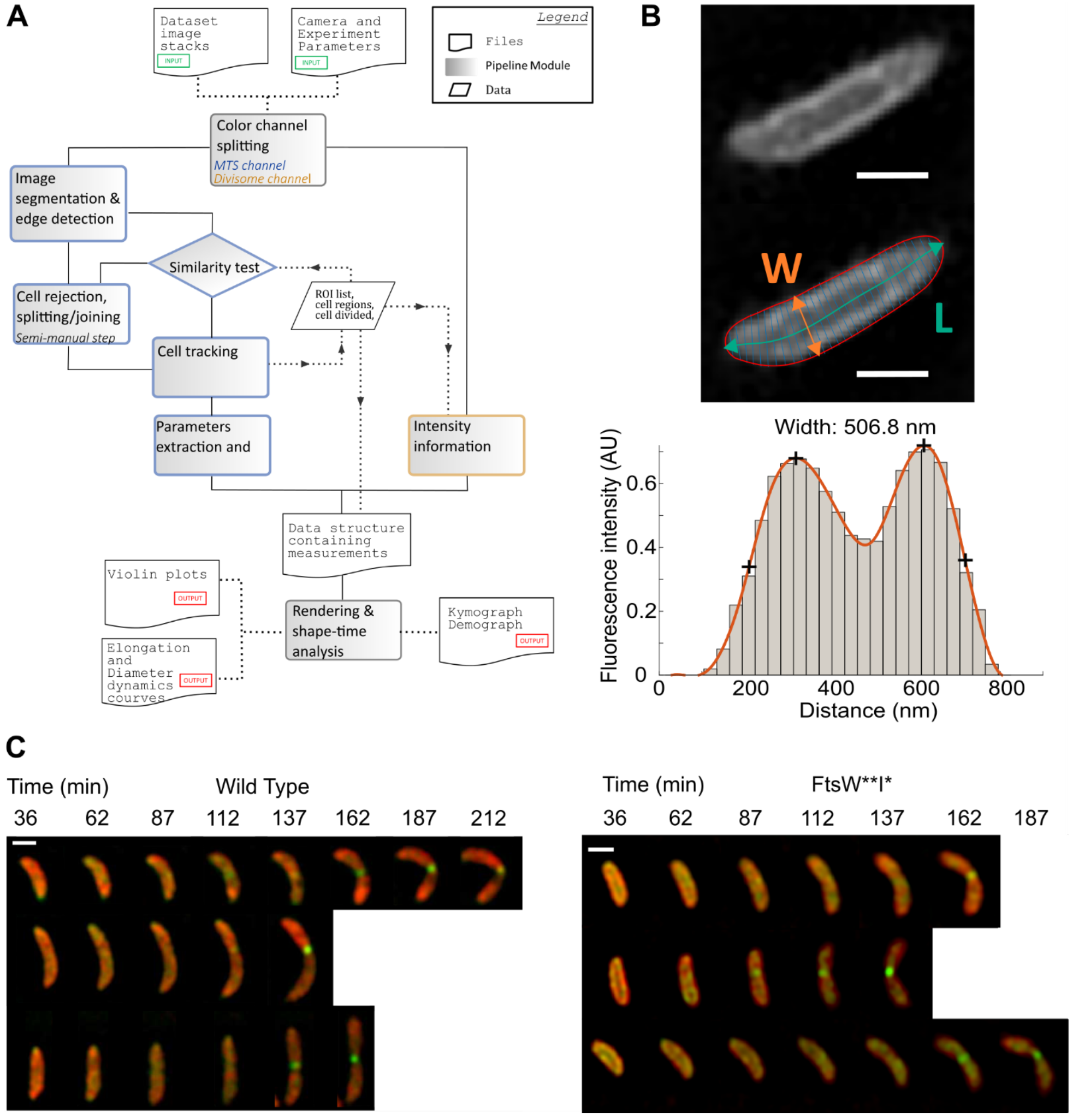
Related to Figure 1 and Methods. Image analysis pipeline. (**A**)Analysis software flowchart. (**B**) Analysis of SIM images inner membrane label: from the raw data, the centerline is calculated. At equally spaced points along the centerline, the width is measured by extracting the intensity profile along a line (thin cyan colored line) perpendicular to the centerline (thick cyan colored line) at that point (top panel red line represent the contour). Lower panel: The extracted intensity profile (grey bars) is smoothed by fitting a spline (orange line lower panel). The maxima are calculated (top plus signs on orange line), and the outer position with the half-maximum value is found (lower plus sign on orange line). The distance between these two positions is defined as the width. (**C**) SlM images of FtsW onset. Red inner membrane MTS2-mCherry labeled, FtsW-GFP label Every one in five frames is shown for three representative cells for Wild Type and FtsW**I*.

**Figure S2.**
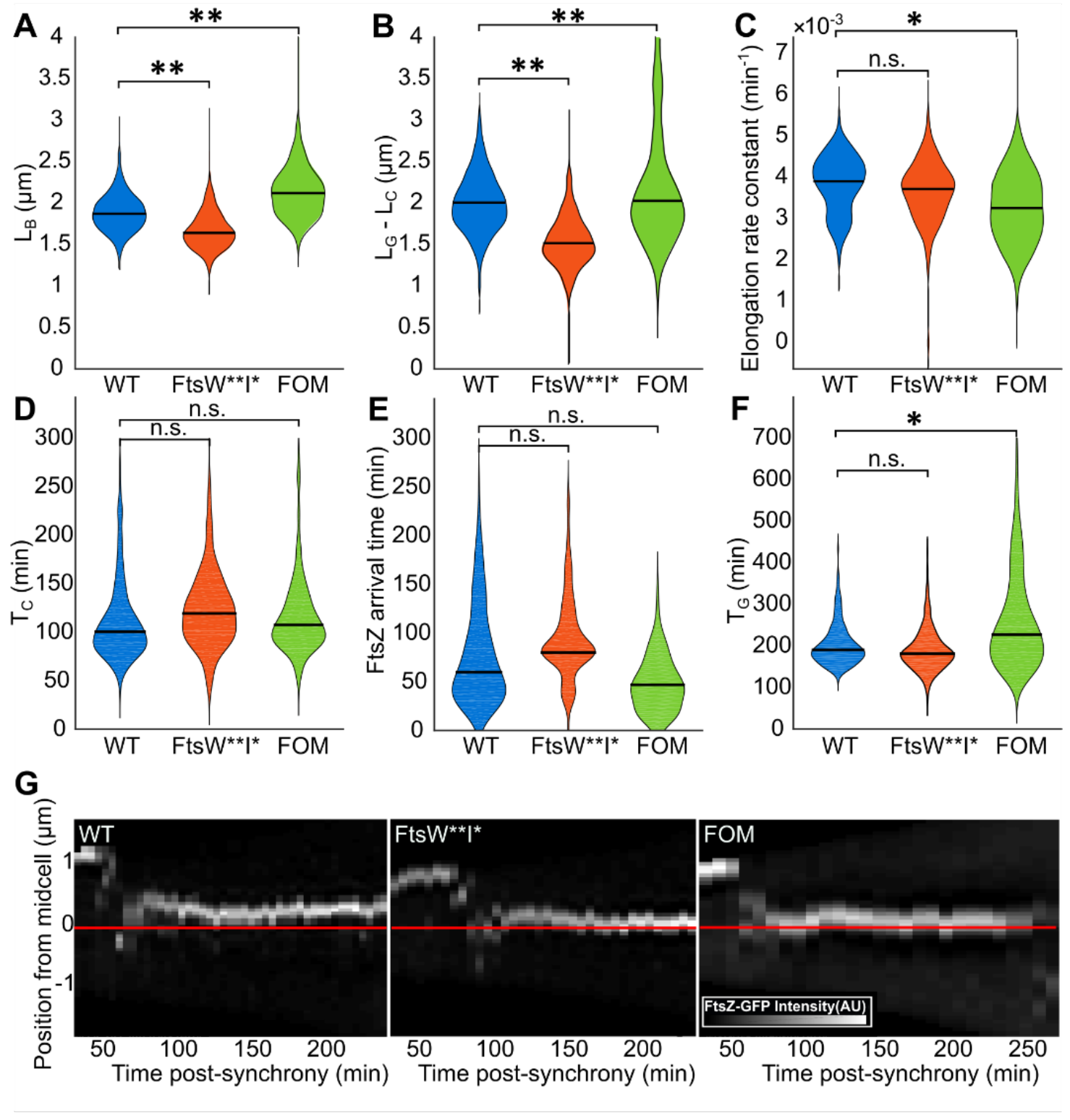
Related to Figure 2. Differences in size and elongation rate between populations. (**A**)Violin plots distributions of length at birth, (**B**) Violin plots distributions of total elongation, (**C**) Violin plots distributions of elongation rate constant, (**D**) Violin plots distributions of onset of constriction, (**E**) Violin plots distributions of FtsZ arrival time and (**F**) Violin plots distributions of generation time for the WT, FtsW**I* and fosfomycin treated WT populations. Black horizontal bars represent the median. N: for (**A-D**, **F**): WT: 406, FtsW**I*: 357, FOM: 203, for (**E**): WT: 207, FtsW**I*: 203, FOM: 105. Significance: **: p<0.005, *: p<0.05, n.s.: not significant. (**G**) Kymographs of representative cells: FtsZ-GFP intensity distribution along the cell’s length (vertical axis), versus cell cycle time (horizontal axis).

**Figure S3.**
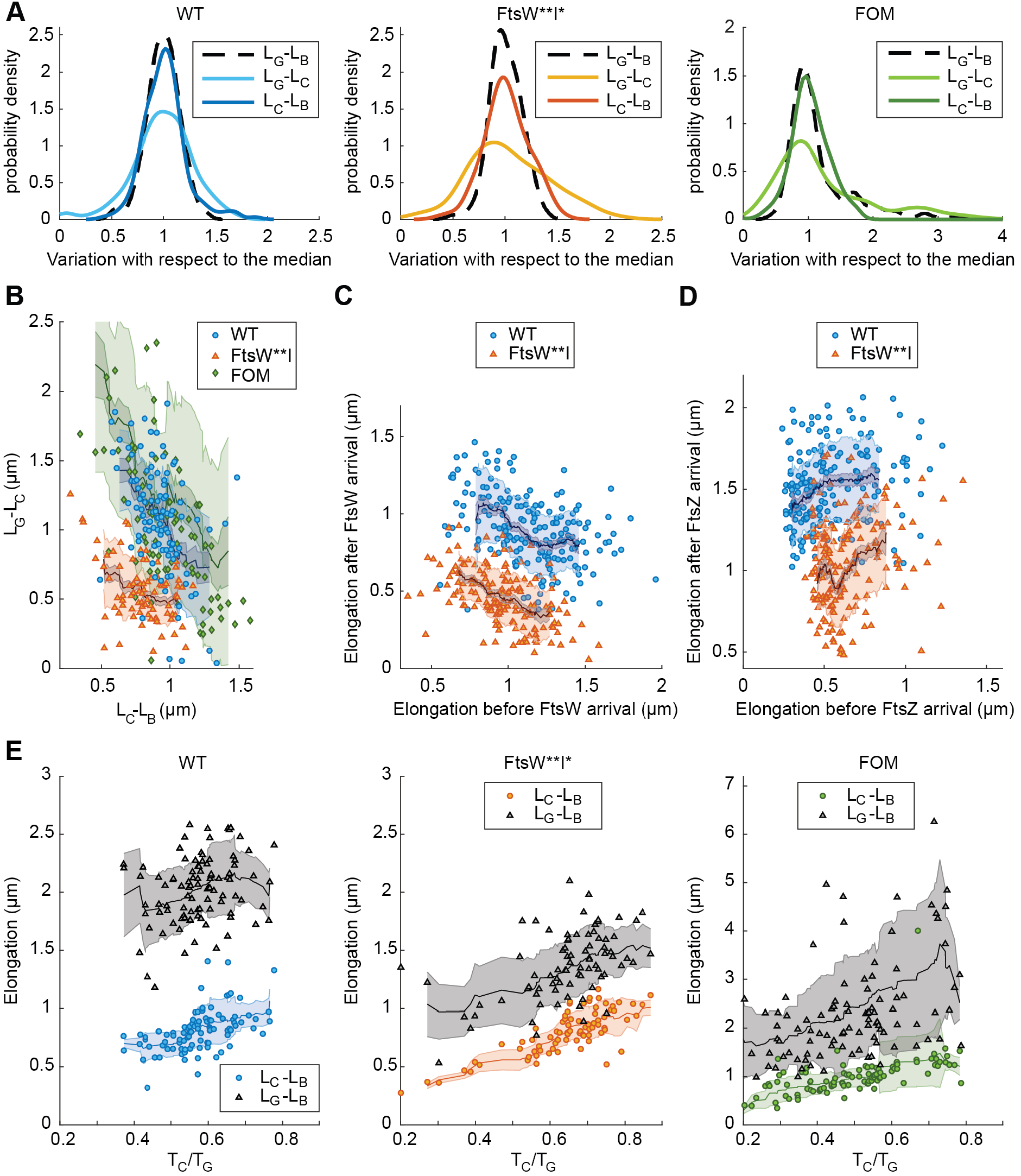
Related to Figure 3. Constriction rate drives compensation between LG-LC and LC-LB. **(A)** Variance in total elongation is smaller than the sum of the variance in elongation before and during constriction. Distribution of the measured elongation divided by median elongation. Onset defined by FtsW arrival. The variance of the sum of independent normal random variable *a* and *b* is equal to the sum of the variance: 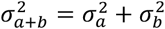 where a and b are the elongation before and during constriction respectively. 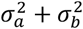 is 0.144 μm^2^ for the WT, 0.080 μm^2^ for FtsW**I* and 1.26 *m^2^ for FOM, which is significantly higher than 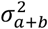 measured from the final total elongation distribution, which is 0.101 μm^2^ for WT, 0.045 μm^2^ for FtsW**I* and 1.04 μm^2^ for FOM. (**B**) Scatterplot showing the elongation during constriction versus the elongation before constriction. Spearman correlation coefficients are: WT:r=−0.44, FtsW**I*: r=−0.32, FOM:r=−0.52, (**C**) Scatterplot of elongation during constriction versus before constriction onset. Spearman correlation: WT: r=-0.44, p-value<0.005, FtsW**I*: r=−0.49, p-value<0.005 (**D**) Scatterplot of elongation after versus before FtsZ arrival. Spearman correlation: WT: Rho=0.20, p<0.005, Mut: r=0.26, p-value<0.005. (**E**) Compensation disappears upon randomization of elongation during constriction. As in Figure **3**, scatterplots of total elongation and elongation before constriction versus normalized onset time for individual Wild Type, FtsW**I* and fosfomycin-treated cells. However, to calculate the total elongation for these plots, the single cell values for elongation during constriction were added randomly to the elongation before constriction of a different cell. Dark lines in (**B-E**) represent the 20 cells moving average; the shaded zones represent the moving standard deviation. Extreme outliers, deviating by more than two standard deviations have been omitted for the calculation of the moving average.

**Figure S4.**
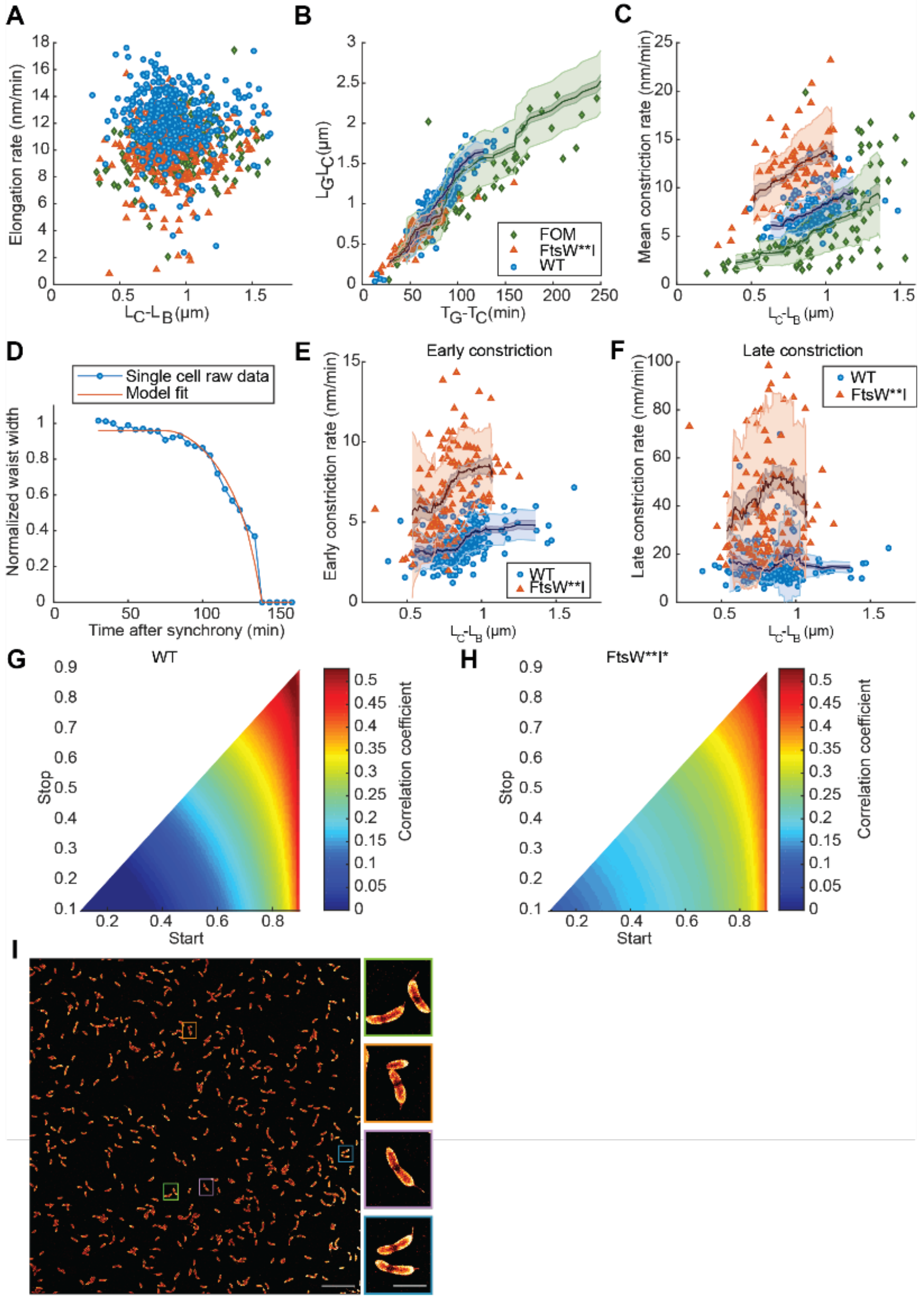
Related to Figure 4. Constriction rate from model fit shows influence of elongation before constriction decreases throughout constriction. (**A**) Elongation rate during constriction is not responsible for compensation: Overall elongation rate during constriction versus elongation rate before constriction. Constriction onset is defined by visible constriction. N and Spearman’s correlation coefficient: WT: N=408, r=-0.11, p-value = 0.026, FtsW**I*: N = 358, r = −0.023, p-value = 0.67, FOM: N = 215, r = 0.30, p-value = 7×10^−6^. (**B**) Scatterplot showing elongation during constriction versus the duration of constriction. Spearman correlation coefficients: WT:r = 0.90, FtsW**I*: r = 0.85, FOM: r = 0.94. N: WT: 96, FtsW**I*: 80, FOM: 102. (**C**) Scatterplot showing the relative duration of constriction ((TG-TC)/TG) versus the elongation before constriction. Spearman Correlation coefficients: WT: r = 0.49 p-value<0.005, FtsW**I*: r = 0.46, p-value<0.005. N: WT: 96, FtsW**I*: 80, FOM: 102. (**D**) Example of a single cell normalized waist width versus time, fitted with the empirical constriction model. (**E**) Early constriction rate, from the fit to the empirical constriction model (supplementary note 2), in function of elongation before constriction. Early constriction rate is defined as the having a normalized waist width between 0.9.and 0.6.(**F**) Late constriction rate, from the fit to the empirical constriction model, in function of elongation before constriction. Late constriction rate is defined as having a waist width between 0.6 and 0.3. (**F**) WT: r = 0.11, p-value=0.13, FtsW**I*: r = 0.20, p-value=0.03. (**G**) Heatmap of correlation coefficient between elongation before constriction and constriction rate, in function of over which portion of constriction the constriction rate is calculated. As in (**E**) and (**F**), the correlation between elongation before constriction and constriction rate was calculated. This was repeated for the constriction rate during various sub-periods of the constriction process, defined by the normalized waist width at the “start” and “stop” of the sub-period. (**H**) Same as (**G**), but for FtsW**I*. Dark lines in (**B, C, E and** F) represent the 20 cells moving average; the shaded zones represent the moving standard deviation. Extreme outliers, deviating by more than two standard deviations have been omitted for the calculation of the moving average (**I**) STORM image of fixed C. crescentus cells stained with fluorescent wheat germ agglutinin. Left panel 100 by 100 microns field of view of fixed C. crescentus cells stained with fluorescent wheat germ agglutinin, Right panels: Magnified views of the bacteria colors correspond to the area selected on the left panel. We used WGA-Alexa647 conjugated dye to stain C. crescentus cell wall. Scale bars 10 μm (Left panel), and 1 μm (Right 4 panels).

## References

1. Koch, A.L. (1996). What size should a bacterium be? A question of scale. Annu. Rev. Microbiol. 50, 317–348.

2. Wang, P., Robert, L., Pelletier, J., Dang, W.L., Taddei, F., Wright, A., and Jun, S. (2010). Robust growth of escherichia coli. Curr. Biol. 20, 1099–1103.

3. National Research Council (US) Steering Group for the Workshop on Size Limits of Very Small Microorganisms (1999). Size Limits of Very Small Microorganisms: Proceedings of a Workshop (Washington (DC): National Academies Press (US)) Available at: http://www.ncbi.nlm.nih.gov/books/NBK224756/ [Accessed August 4, 2017].

4. Donachie, W.D., and Begg, K.J. (1989). Cell length, nucleoid separation, and cell division of rod shaped and spherical cells of Escherichia coli. J. Bacteriol. 171, 4633–4639.

5. Beveridge, T.J. (1988). The bacterial surface: General considerations towards design and function. Can. J. Microbiol. 34, 363–372.

6. Schulz, H.N., and Jørgensen, B.B. (2001). Big bacteria. Annu. Rev. Microbiol. 55, 105–137.

7. Marczynski, G.T. (1999). Chromosome Methylation and Measurement of Faithful, Once and Only Once per Cell Cycle Chromosome Replication inCaulobacter crescentus. J. Bacteriol. 181, 1984–1993.

8. Cooper, S., and Helmstetter, C.E. (1968). Chromosome replication and the division cycle of Escherichia coli Br. J. Mol. Biol. 31, 519–540.

9. Skerker, J.M., and Laub, M.T. (2004). Cell-cycle progression and the generation of asymmetry in Caulobacter crescentus. Nat. Rev. Microbiol. 2, 325–337.

10. Aaron, M., Charbon, G., Lam, H., Schwarz, H., Vollmer, W., and Jacobs-Wagner, C. (2007). The tubulin homologue FtsZ contributes to cell elongation by guiding cell wall precursor synthesis in Caulobacter crescentus. Mol. Microbiol. 64, 938–952.

11. Kuru, E., Hughes, H.V., Brown, P.J., Hall, E., Tekkam, S., Cava, F., De, P., Brun, Y.V., and Vannieuwenhze, M.S. (2012). In situ probing of newly synthesized peptidoglycan in live bacteria with fluorescent D-amino acids. Angew. Chem.-Int. Ed. 51, 12519–12523.

12. Taheri-Araghi, S., Bradde, S., Sauls, J.T., Hill, N.S., Levin, P.A., Paulsson, J., Vergassola, M., and Jun, S. (2015). Cell-Size Control and Homeostasis in Bacteria. Curr. Biol. 25, 385–391.

13. Weart, R.B., Lee, A.H., Chien, A.-C., Haeusser, D.P., Hill, N.S., and Levin, P.A. (2007). A metabolic sensor governing cell size in bacteria. Cell 130, 335–347.

14. Coltharp, C., Buss, J., Plumer, T.M., and Xiao, J. (2016). Defining the rate-limiting processes of bacterial cytokinesis. Proc. Natl. Acad. Sci. U. S. A. 113, E1044–E1053.

15. Thanbichler, M., and Shapiro, L. (2006). MipZ, a Spatial Regulator Coordinating Chromosome Segregation with Cell Division in Caulobacter. Cell 126, 147–162.

16. Wallden, M., Fange, D., Lundius, E.G., Baltekin, O., and Elf, J. (2016). The Synchronization of Replication and Division Cycles in Individual E. coli Cells. Cell 166, 729–739.

17. Sompayrac, L., and Maaløe, O. (1973). Autorepressor Model for Control of DNA Replication. Nature. New Biol. 241, 133.

18. Harris, L.K., and Theriot, J.A. (2016). Relative rates of surface and volume synthesis set bacterial cell size. Cell 165, 1479–1492.

19. Campos, M., Surovtsev, I.V., Kato, S., Paintdakhi, A., Beltran, B., Ebmeier, S.E., and Jacobs-Wagner, C. (2014). A constant size extension drives bacterial cell size homeostasis. Cell 159, 1433–1446.

20. Banerjee, S., Lo, K., Daddysman, M.K., Selewa, A., Kuntz, T., Dinner, A.R., and Scherer, N.F. (2017). Biphasic growth dynamics control cell division in Caulobacter crescentus. Nat. Microbiol. 2, 17116.

21. Jun, S., and Taheri-Araghi, S. (2015). Cell-size maintenance: universal strategy revealed. Trends Microbiol. 23, 4–6.

22. den Blauen, T., Hamoen, L.W., and Levin, P.A. (2017). The divisome at 25: the road ahead. Curr. Opin. Microbiol. 36, 85–94.

23. Leitao, R.M., and Kellogg, D.R. (2017). The duration of mitosis and daughter cell size are modulated by nutrients in budding yeast. J Cell Biol, jcb.201609114.

24. Gustafsson, M.G.L. (2000). Surpassing the lateral resolution limit by a factor of two using structured illumination microscopy. J. Microsc. 198, 82–87.

25. Kahan, F.M., Kahan, J.S., Cassidy, P.J., and Kropp, H. (1974). THE MECHANISM OF ACTION OF FOSFOMYCIN (PHOSPHONOMYCIN). Ann. N. Y. Acad. Sci. 235, 364–386.

26. Meeske, A.J., Riley, E.P., Robins, W.P., Uehara, T., Mekalanos, J.J., Kahne, D., Walker, S., Kruse, A.C., Bernhardt, T.G., and Rudner, D.Z. (2016). SEDS proteins are a widespread family of bacterial cell wall polymerases. Nature 537, 634–638.

27. Adam, M., Fraipont, C., Rhazi, N., Nguyen-Distèche, M., Lakaye, B., Frère, J.M., Devreese, B., Beeumen, J.V., Heijenoort, Y., van, Heijenoort, J. van, et al. (1997). The bimodular G57-V577 polypeptide chain of the class B penicillin-binding protein 3 of Escherichia coli catalyzes peptide bond formation from thiolesters and does not catalyze glycan chain polymerization from the lipid II intermediate. J. Bacteriol. 179, 6005–6009.

28. Modell, J.W., Kambara, T.K., Perchuk, B.S., and Laub, M.T. (2014). A DNA Damage-Induced, SOS-Independent Checkpoint Regulates Cell Division in Caulobacter crescentus. PLoS Biol. 12.

29. Sargent, M.G. (1975). Control of cell length in Bacillus subtilis. J. Bacteriol. 123, 7–19.

30. Schaechter, M., Maaloe, O., and Kjeldgaard, N.O. (1958). Dependency on medium and temperature of cell size and chemical composition during balanced grown of Salmonella typhimurium. J. Gen. Microbiol. 19, 592–606.

31. Beaufay, F., Coppine, J., Mayard, A., Laloux, G., Bolle, X.D., and Hallez, R. (2015). A NAD-dependent glutamate dehydrogenase coordinates metabolism with cell division in Caulobacter crescentus. EMBO J. 34, 1786–1800.

32. Morales Angeles, D., Liu, Y., Hartman, A.M., Borisova, M., de Sousa Borges, A., de Kok, N., Beilharz, K., Veening, J.-W., Mayer, C., Hirsch, A.K.H., et al. (2017). Pentapeptide-rich peptidoglycan at the Bacillus subtilis cell-division site. Mol. Microbiol. 104, 319–333.

33. Yahashiri, A., Jorgenson, M.A., and Weiss, D.S. (2015). Bacterial SPOR domains are recruited to septal peptidoglycan by binding to glycan strands that lack stem peptides. Proc. Natl. Acad. Sci. U. S. A. 112, 11347–11352.

34. Goley, E.D., Comolli, L.R., Fero, K.E., Downing, K.H., and Shapiro, L. (2010). DipM links peptidoglycan remodeling to outer membrane organization in Caulobacter. Mol. Microbiol. 77, 56–73.

35. Möll, A., Schlimpert, S., Briegel, A., Jensen, G.J., and Thanbichler, M. (2010). DipM, a new factor required for peptidoglycan remodelling during cell division in Caulobacter crescentus. Mol. Microbiol. 77, 90–107.

36. Poggio, S., Takacs, C.N., Vollmer, W., and Jacobs-Wagner, C. (2010). A protein critical for cell constriction in the Gram-negative bacterium Caulobacter crescentus localizes at the division site through its peptidoglycan-binding LysM domains. Mol. Microbiol. 77, 74–89.

37. Douglass, K.M., Sieben, C., Archetti, A., Lambert, A., and Manley, S. (2016). Super-resolution imaging of multiple cells by optimized flat-field epi-illumination. Nat. Photonics 10, 705–708.

38. Bisson-Filho, A.W., Hsu, Y.-P., Squyres, G.R., Kuru, E., Wu, F., Jukes, C., Sun, Y., Dekker, C., Holden, S., VanNieuwenhze, M.S., et al. (2017). Treadmilling by FtsZ filaments drives peptidoglycan synthesis and bacterial cell division. Science 355, 739–743.

39. Yang, X., Lyu, Z., Miguel, A., McQuillen, R., Huang, K.C., and Xiao, J. (2017). GTPase activity-coupled treadmilling of the bacterial tubulin FtsZ organizes septal cell wall synthesis. Science 355, 744–747.

40. Goley, E.D., Yeh, Y.-C., Hong, S.-H., Fero, M.J., Abeliuk, E., McAdams, H.H., and Shapiro, L. (2011). Assembly of the Caulobacter cell division machine. Mol. Microbiol. 80, 1680–1698.

41. Holden, S.J., Pengo, T., Meibom, K.L., Fernandez, C.F., Collier, J., and Manley, S. (2014). High throughput 3D super-resolution microscopy reveals Caulobacter crescentus in vivo Z-ring organization. Proc. Natl. Acad. Sci. 111, 4566–4571.

42. Schrader, J.M., and Shapiro, L. (2015). Synchronization of Caulobacter Crescentus for Investigation of the Bacterial Cell Cycle. JoVE J. Vis. Exp., e52633–e52633.

43. Evinger, M., and Agabian, N. (1977). Envelope-associated nucleoid from Caulobacter crescentus stalked and swarmer cells. J. Bacteriol. 132, 294–301.

44. Thanbichler, M., Iniesta, A.A., and Shapiro, L. (2007). A comprehensive set of plasmids for vanillate-and xylose-inducible gene expression in Caulobacter crescentus. Nucleic Acids Res. 35, e137.

45. Szeto, T.H., Rowland, S.L., Habrukowich, C.L., and King, G.F. (2003). The MinD Membrane Targeting Sequence Is a Transplantable Lipid-binding Helix. J. Biol. Chem. 278, 40050–40056.

46. Pincus, Z., and Theriot, J.A. (2007). Comparison of quantitative methods for cell-shape analysis. J. Microsc. 227, 140–156.

47. Schneider, C.A., Rasband, W.S., and Eliceiri, K.W. (2012). NIH Image to ImageJ: 25 years of image analysis. Nat. Methods 9, 671–675.

48. Sliusarenko, O., Heinritz, J., Emonet, T., and Jacobs-Wagner, C. (2011). High-throughput, subpixel precision analysis of bacterial morphogenesis and intracellular spatio-temporal dynamics. Mol. Microbiol. 80, 612–627.

## References

1. Preibisch, S., Saalfeld, S., Schindelin, J., and Tomancak, P. (2010). Software for bead-based registration of selective plane illumination microscopy data. Nat. Methods 7, 418–419.

2. Szeto, T.H., Rowland, S.L., Habrukowich, C.L., and King, G.F. (2003). The MinD Membrane Targeting Sequence Is a Transplantable Lipid-binding Helix. J. Biol. Chem. 278, 40050–40056.

3. Nobuyuki Otsu (1979). A Threshold Selection Method from Gray-Level Histograms. IEEE Trans. SYSTREMS MAN Cybern. SMC-9,.

4. Sliusarenko, O., Heinritz, J., Emonet, T., and Jacobs-Wagner, C. (2011). High-throughput, subpixel precision analysis of bacterial morphogenesis and intracellular spatio-temporal dynamics. Mol. Microbiol. 80, 612–627.

5. Pincus, Z., and Theriot, J.A. (2007). Comparison of quantitative methods for cell-shape analysis. J. Microsc. 227, 140–156.

6. Sycuro, L.K., Pincus, Z., Gutierrez, K.D., Biboy, J., Stern, C.A., Vollmer, W., and Salama, N.R. (2010). Peptidoglycan crosslinking relaxation promotes Helicobacter pylori’s helical shape and stomach colonization. Cell 141, 822–833.

7. Harris, L.K., and Theriot, J.A. (2016). Relative Rates of Surface and Volume Synthesis Set Bacterial Cell Size. Cell 165, 1479–1492.

8. Coltharp, C., Buss, J., Plumer, T.M., and Xiao, J. (2016). Defining the rate-limiting processes of bacterial cytokinesis. Proc. Natl. Acad. Sci. U. S. A. 113, E1044–1053.

